# Modeling Combined Ultrasound and Photoacoustic Imaging: Simulations aiding Device Development and Deep Learning

**DOI:** 10.1101/2020.11.07.371930

**Authors:** Sumit Agrawal, Ajay Dangi, Sri-Rajasekhar Kothapalli

## Abstract

Combined ultrasound and photoacoustic (USPA) imaging has attracted several clinical applications due to its ability to simultaneously display structural and molecular information of deep biological tissue in real time. However, the depth dependent optical attenuation and the unknown optical and acoustic heterogeneities, limit the USPA imaging performance, especially from deeper tissue regions. Novel instrumentation, image reconstruction and deep learning methods are currently being explored to improve the USPA image quality. Effective implementation of these approaches requires a reliable USPA simulation tool capable of generating US based anatomical and PA based molecular information. Here, we developed a hybrid USPA simulation platform by integrating finite element models of light and ultrasound propagation. The feasibility of modeling US combined with optical fluence dependent multispectral PA imaging is demonstrated using in silico homogeneous and heterogeneous prostate tissue. The platform allows optimization of device design parameters, such as the aperture size and frequency of light source and ultrasound detector arrays. In addition, the potential of this simulation platform to generative massive USPA datasets aiding the data driven deep-learning enhanced USPA imaging has been validated using both simulations and experiments.

## I. Introduction

**P**hotoacoustic imaging (PAI) gained significant attention of biomedical research community by displaying rich molecular optical absorption contrast images of deep tissue with higher spatial resolution than possible with pure optical imaging technologies [1]. In PAI, the light absorbing tissue chromophores (e.g., hemoglobin and melanin) undergo thermoelastic expansion and generate broadband ultrasound waves, i.e. photoacoustic waves. These waves propagate and get detected by the ultrasound (US) transducers placed outside the body, to form 3-D photoacoustic (PA) images with rich molecular contrast. In PAI, the spatial resolution and the penetration depth in the photon diffusion regime (> mm) are scalable with ultrasound frequency, and the temporal resolution is mostly limited by the laser pulse-repetition frequency. Multi-wavelength PAI further provides functional (e.g., oxygen saturation) and molecular-specific (e.g., melanin) information important to diagnose the diseased tissue [2], [3]. The ease of integration of PAI capabilities to the clinical US systems further allowed the demonstration of dual-modality USPA devices capable of simultaneously displaying in real-time the anatomical and molecular information in pre-clinical as well as clinical research [4], [5]. In recent years, many clinical applications of handheld USPA devices have emerged; such as imaging vasculature of human breast [6–8], prostate [9], ovaries [10], muscular dystrophy [11], melanoma metastasis [12], inflammatory arthritis [13], thyroid [14] and Crohn’s disease [15].

However, the USPA imaging commonly suffers from depth dependent optical and acoustic attenuation which affects the visibility of deep tissue targets [16], [17]. Moreover, the unknown optical and acoustic heterogeneities complicate the estimation of optical fluence needed for quantitative PAI [18]. These heterogeneities also introduce reflection artifacts [19]. Together, these factors limit the USPA imaging performance, especially for deep tissue clinical applications.

To overcome these limitations, novel USPA imaging hardware [20–24], image reconstruction [25], deep learning and artifact correction methods [26], [27] are actively being investigated. These approaches would benefit from accurate simulation of USPA devices while taking into account the optical and acoustic parameters of the device and the biological tissue of interest. Computational models reduce the total cost and time involved in the device development and validations through phantom, animal and clinical studies. The credibility of such simulation models needs to be assessed through the extensive verification and validation studies against the experiments. Towards this goal, several PAI simulation approaches have been proposed to optimize the PAI device geometries and quantitative image reconstruction [28–30]. These simulations have been predominantly reported for photoacoustic computed tomography geometries that involve rotating single element transducers or sparsely distributed ultrasound transducer elements [28]. However, despite rapid progress in the development and clinical translation of dual-modality USPA devices, no simulation studies have been reported to the best of our knowledge for modeling dual-modality B-mode US and PA imaging of realistic heterogeneous tissue medium. Recently, PAI simulations using K-Wave toolbox have become popular [31]. However, K-Wave does not account for the realistic PAI scenarios where the optical fluence strongly depends on the heterogeneous optical properties of the tissue, irradiation wavelength and attenuates as a function of tissue depth. Nima et al. [29] recently used Monte Carlo simulations for estimating optical fluence in a homogeneous tissue phantom and reconstructed PA images using K-Wave toolbox. However, this study did not report B-mode US images to display corresponding structural information of the tissue phantom. In addition, no multispectral PAI results were reported to delineate the molecular information of the tissue. Moreover, Monte Carlo simulations are computationally expensive and slow. Overcoming these challenges, we recently reported a finite-element model (NIRFast toolbox [32]) that uses diffusion approximation for estimating the optical fluence in a homogeneous medium [33]. This depth dependent optical fluence was then converted to initial pressure rise and back-propagated using K-Wave toolbox to obtain simulated B-mode PA images. Expanding on this conference proceedings paper, here we present a dual-modality USPA simulation platform that displays US based anatomical and PA based molecular information in a realistic tissue background. To assess the credibility of our simulation models, we performed parametric verification and validation studies against the experiments. In addition, we demonstrate the applicability of the USPA simulation datasets aiding the data driven deep learning algorithms to locate deep tissue PAI targets in high scattering noise and effectively unmixing the molecular information.

The rest of the paper is organized as follows: Section II presents the simulation geometry, including the ultrasound and optical properties of the tissue phantoms, and the workflow of the hybrid USPA imaging simulations. Section III presents parametric verification and validation study results and discussions. Section IV concludes the work with insights to the future scope of the work.

## II. Methodology

In this section, we start with the description of simulation geometry and different device configurations. Later we describe the optical properties of the medium and the light absorbing (target) objects used for the simulations presented in this work. A schematic representation and workflow of our simulation platform for generating dual-modality US as well as PA images is also presented. The bulk optical properties of the medium and associated photon transport mean free path determines the fraction of photons (optical fluence) present at each location, and was achieved using an open-source software package, NIRFast toolbox [32]. The fluence map is then converted to the pressure distribution, which is then propagated and detected using an open source MATLAB based software platform, K-wave toolbox [31], In contrast, the simulation of B-mode ultrasound image formation is rather straightforward using K-Wave toolbox. Here, ultrasound imaging is performed using the same ultrasound transducer array that is used for photoacoustic detection.

### A. Simulation Geometry

Here we introduce the phantom geometry, and key design parameters for light source and ultrasound transducer arrays employed in the USPA imaging device modeled in our simulations. Fig. 1 shows three device configurations used for studying the effect of the aperture size and frequency of light source as well as ultrasound transducer on the USPA imaging performance. The phantom geometry is designed as a rectangular two-dimensional (2D) grid of 100 mm width and 60 mm depth. The grid spacing is 0.2 mm and the total number of nodes in the grid are 150,801. The linear ultrasound transducer and light source arrays are both kept at the bottom of the tissue grid, centered at the (x, z) = (0, 0) origin of the grid. In all simulations, the ultrasound transducer array is centered at 1 MHz frequency with a fractional bandwidth of 80%, unless otherwise specified. Fig. 1(a) shows the first USPA device configuration consisting of a 64-element ultrasound transducer array spread from −6.4 mm to 6.4 mm (total of 12.8 mm), and a 40 mm long light source from −20 mm to 20 mm, both along the x-direction. Fig. 1(b) shows the second device configuration consisting of a 128-element transducer array spread from −12.8 mm to 12.8 mm (total of 25.6 mm), and a 40 mm long light source. Fig. 1(c) shows the third device with 20 mm long light source and a 128-element transducer array.

**Fig. 1.**
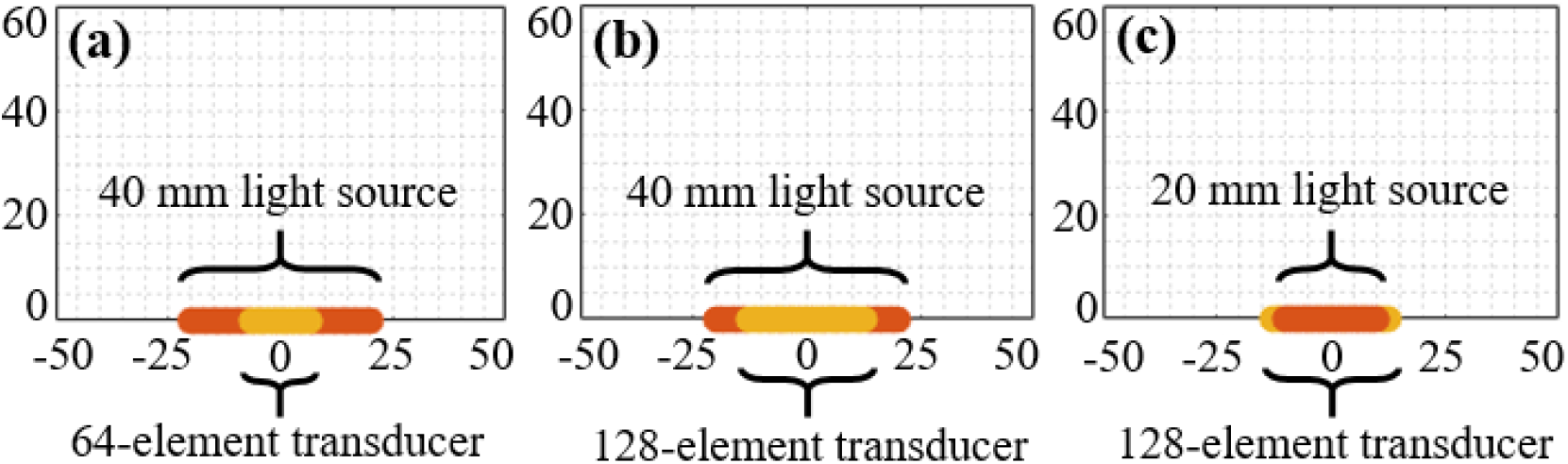
Phantom geometries for three USPA device configurations used in parametric simulation studies with varying ultrasound transducer array and light apertures. Transducer array: yellow; light source: orange; scale: mm.

### B. Optical Properties of tissue mimicking phantom

The optical properties of the tissue phantoms determine the amount of light absorbed or scattered at each grid position. In our simulations, we have used both homogenous as well as heterogeneous backgrounds mimicking human prostate tissue. The light absorbing molecules such as oxy-hemoglobin (HbO_2_), deoxy-hemoglobin (Hb), and exogenous contrast agent indocyanogreen (ICG) are used as molecular targets placed in the tissue background. In Table I, we list the absorption coefficients of the three molecular targets and the tissue background, taken from [34] and [35]. The reduced scattering coefficients (*μ′s*) for the tissue background and the molecular targets (HbO_2_, Hb, and ICG) are calculated using Eq. 1:

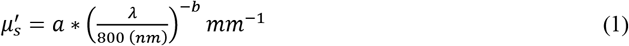

where, *a* (mm^-1^) is the value of reduced scattering coefficient at 800 nm, *λ* is the wavelength at which the scattering coefficient is being calculated, and *b* is the scattering power [34].

**TABLE I:** Absorption and reduced scattering coefficients (in mm^-1^) of molecular targets (HbO_2_, Hb, and ICG), background tissue used in simulations.

| $\lambda$<br>(nm) | $\text{HbO}_2$ | | $\text{Hb}$ | | ICG | | Background | |
| --- | --- | --- | --- | --- | --- | --- | --- | --- |
| | $\mu_a$ | $\mu'_s$ | $\mu_a$ | $\mu'_s$ | $\mu_a$ | $\mu'_s$ | $\mu_a$ | $\mu'_s$ |
| 750 | 0.26 | 1.04E-5 | 0.78 | 1.04E-5 | 0.0982 | 1.04E-5 | 0.0013 | 1.1368 |
| 775 | 0.34 | 1.02E-5 | 0.62 | 1.02E-5 | 0.1654 | 1.02E-5 | 0.0011 | 1.0805 |
| 800 | 0.425 | 1.00E-5 | 0.420 | 1.00E-5 | 0.1257 | 1.00E-5 | 0.0011 | 1.0287 |
| 825 | 0.475 | 9.79E-5 | 0.376 | 9.79E-5 | 0.0258 | 9.79E-5 | 0.0013 | 0.9808 |
| 850 | 0.53 | 9.60E-5 | 0.376 | 9.60E-5 | 0.0067 | 9.60E-5 | 0.0013 | 0.936 |
| 875 | 0.593 | 9.42E-5 | 0.376 | 9.42E-5 | 0.0003 | 9.42E-5 | 0.0018 | 0.8953 |
| 900 | 0.625 | 9.25E-5 | 0.421 | 9.25E-5 | 0 | 9.25E-5 | 0.0041 | 0.8571 |

### C. Fluence Calculation

The first step in the generation of PA image involves the forward propagation of light into the tissue phantom and calculation of optical fluence. We have used an open-source software package, NIRFast [32] that solves for the light diffusion equation and calculates the light fluence, *ϕ* at each grid position of the 2D phantom. Fig. 2 shows that the optical fluence reaches deeper tissue depths for a light aperture size of 40 mm (Fig. 2a) as compared to 20 mm (Fig. 2b), inside a homogeneous tissue background described in Table I at 800 nm wavelength. Fig. 2c plots the wavelength-dependent optical fluence attenuation as a function of depth inside the homogeneous tissue background calculated for a 40 mm light aperture size. These plots were used for compensating the wavelength dependent fluence attenuation as a function of depth in all our photoacoustic imaging simulations [36].

**Fig. 2.**
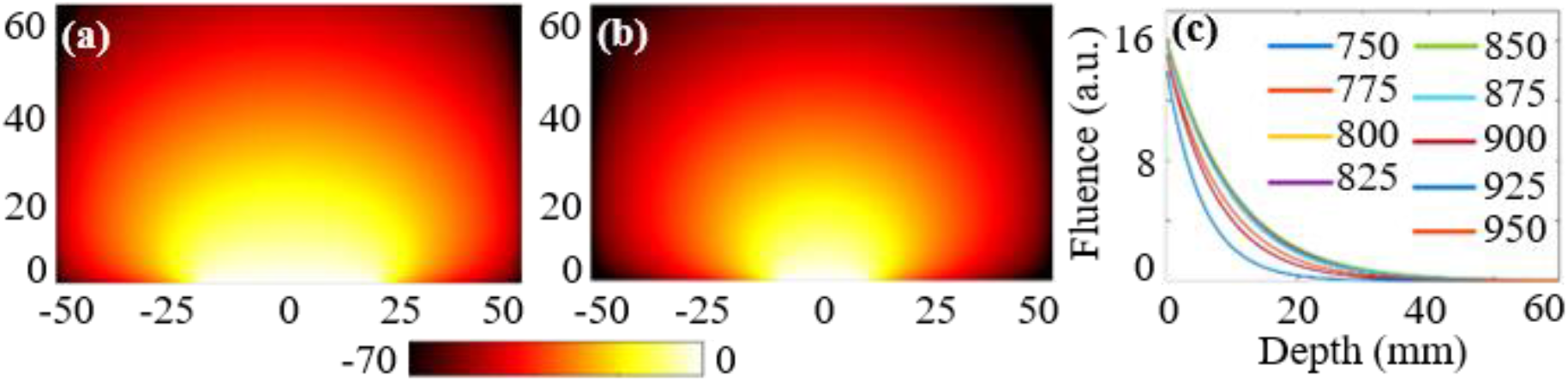
optical fluence calculations inside the tissue medium. (a, b) Fluence maps generated by a 40 mm, and a 20 mm aperture size light source, respectively. (c) Wavelength dependent attenuation of optical fluence as a function of depth. Scale: mm, colorbar: dB.

### D. Flowchart for US and PA image generation

This section presents the systematic workflow (Fig. 3) of the USPA simulation platform for generating US and PA images. We designed a combined acoustic and optical phantom consisting of nine circular targets of radius 0.25 mm, shown in Fig. 3a. For the US phantom, the acoustic impedances are set to 1.5 and 1.7 MRayl for the background and the circular targets, respectively [37]. For the optical phantom, six targets out of nine (2, 3, 5, 6, 8, and 9 marked with black circles) are defined as light absorbing vascular targets having a higher absorption coefficient of 0.425 mm^-1^ (typical blood absorption at 800 nm) compared to the other three targets (1, 4, and 7 marked with orange circles) having an absorption coefficient of 0.0011 mm^-1^ matching with the tissue background [34].

**Fig. 3.**
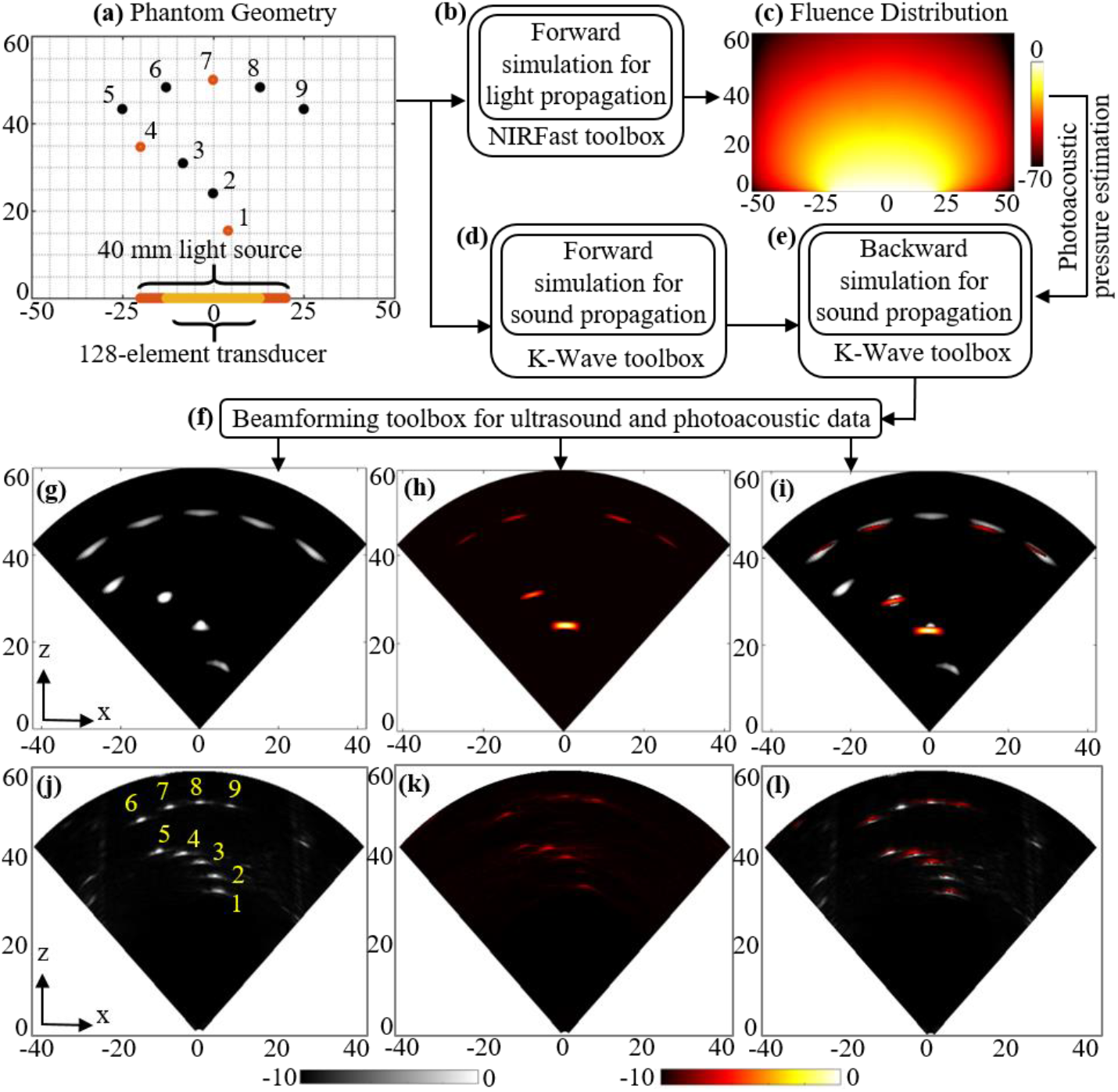
(a-f) Flowchart presenting key steps involved in the USPA simulations. US, PA and coregistered US+PA images of a homogeneous phantom consisting of nine-circular targets obtained from (g-i) simulations, and (j-l) experiments. Scale: mm, colorbar: dB.

For the US image generation, the forward and backward simulations involve the propagation of ultrasound waves from the transducer elements to the tissue medium and vice versa, respectively using an open source ultrasound simulation platform K-Wave toolbox [31] (Figs. 3 (d, e)). The received time-dependent pressure data at each transducer location is then fed to our custom developed US beamforming toolbox (Fig. 3f), which uses standard Delay-and-Sum [9] approach with fixed transmit and dynamic receive focusing, to reconstruct B-mode US images, as shown in Fig. 3g.

The first simulation step for a PA image generation involves the forward propagation of light in the discretized 2D optical phantom grid with pre-defined optical properties using NIRfast [32] (Fig. 3b). It uses a finite element method to solve for the radiative transport equation with a diffusion approximation and calculates the light fluence, *ϕ(r,λ)*, at each grid position. This generates the 2D optical fluence map, as shown in Fig. 3c, for a given wavelength (e.g., 800 nm in this case). The fluence map is then converted to the initial pressure map, *P_s_(r, λ)*, using Eq. 2, where *Γ* is the Gruneisen parameter, a measure of conversion efficiency from the light absorption to pressure. A fixed homogeneous value of 0.2 for *Γ* was assumed in all our studies [38]. *ϕ(r, λ)* and *μ_a_(r, λ)* are the optical fluence and the absorption coefficients at a position r and for wavelength *λ* [39].

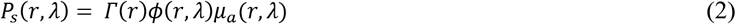

This equation provides the value of initial pressure at each grid location of the tissue phantom. The detection path for PA simulation involves the propagation of generated photoacoustic pressure waves from the respective grid positions inside the tissue phantom to the position of all ultrasound transducer elements using K-Wave toolbox [31] (Fig. 3e). This solves the solution of following acoustic wave equation [39]:

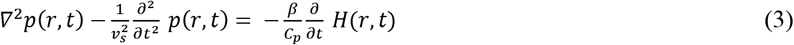

where, *v_s_* is the acoustic speed in the medium, *β* is the isobaric volume expansion coefficient, *C_p_* is the isobaric specific heat, and *H(r, t)* is the amount of thermal energy converted to pressure at a position *r* and time *t*. The pressure data from each grid position is measured as a function of time at each transducer element location in the form of time-dependent sensor voltage data; also termed as radio frequency (RF) channel data. We then use our custom developed receive-only Delay-and-Sum beamforming toolbox (Fig. 3f) on the RF channel data [9] to reconstruct B-mode PA image, as shown in Fig. 3h. Due to the difference in acoustic impedance between each circular target and the background, all nine targets generated ultrasound echoes and are imaged in the US mode (Fig. 3g), whereas, only the six light absorbing vascular targets (represented as black dots, simulating blood vessels) are detected in the PA image (Fig. 3h). In addition to US and PA images, the simulation platform also displays the co-registered US+PA image (Fig. 3i).

In our simulation studies, the speed of sound in homogeneous tissue background was assumed to be 1480 m/s and the frequency (*f*) dependent acoustic attenuation inside the tissue was taken as 0.75 *f*^1.5^ dB/cm/MHz^1.5^ [37–39]. A voxel size of 20 was considered as perfectly matched layer surrounding the tissue grid. In PAI simulations, the fluence map generation (using NIRFast) over the defined 200 μm resolution grid took ~5 minutes and the pressure propagation to gather the raw photoacoustic signals (using K-Wave) took ~30 seconds. On the other hand, US simulations, including both forward excitation and ultrasonic detection over the same 200 μm grid (using K-Wave) took ~2 minutes for every A-line and a total of 101 scan lines were used for acquiring one B-mode US image. All computations were performed over a Xeon processor with 128 GB RAM and 16 GB NVIDIA Titan XP GPU.

To substantiate the imaging capabilities of our simulations, we performed experiments on a tissue mimicking homogeneous intralipid phantom using a custom designed USPA device. The device integrates a linear 64-element CMUT array with a fiber optic light guide connected to a tunable OPO laser (Opotek Inc., 10-Hz repetition rate, 5-ns pulse width, 680- to 950-nm wavelength range). Similar to the simulated phantom, the experimental phantom consisted of nine fishing-wire targets (mimicking the simulated circular targets) in which seven are painted black to mimic the light absorbing vascular targets. The USPA data is acquired and reconstructed using a PC-based multi-channel US data acquisition system (Verasonics, Inc.) [9]. Beamformed US, PA (at 800-nm) and coregistered US+PA images are shown in Figs. 3(j-l). These results confirm that our simulation platform is capable of closely modeling the USPA devices in realistic experimental conditions.

The PA simulation for a single wavelength described in the flowchart can be extended to generate multi-wavelength PA images for obtaining the spectrally unmixed images of different molecular targets, as demonstrated in the subsequent sections.

## III. Results and discussions

This section presents validation experiments conducted to further evaluate the performance of the hybrid USPA simulation platform. Sections III-(A, B) presents the parametric studies such as the effect of aperture sizes of the ultrasound transducer and the light source, and the center frequency of the ultrasound transducer on the USPA imaging performance. Section III-C presents the multispectral PA imaging capabilities of the platform for delineating different molecular targets. Section III-D demonstrates the USPA imaging simulations of heterogeneous prostate tissue. Section III-E presents the feasibility of the simulation platform in aiding deep-learning enhanced PAI.

### A. Effect of ultrasound transducer and light array size

With a fixed 40 mm aperture size of the light source, we studied here the effect of US transducer array size by employing i) a 64-element linear US array of 12.8 mm length (Fig. 1a), and ii) a 128-element linear US array of 25.6 mm length (Fig. 1b), with 0.2 mm pitch. We used the same tissue phantom with nine circular targets as described in section II-D. Figs. 4(a-c) and Figs. 4(d-f) show US, PA, and coregistered US+PA simulated imaging results for the 64-element and 128-element transducer sizes respectively. Quantified spatial resolutions of targets #2 and #8, provided in Table II, demonstrate that the lateral resolution and the visibility of deeper targets (depth of imaging) improves with the aperture size of US transducer.

**Fig. 4.**
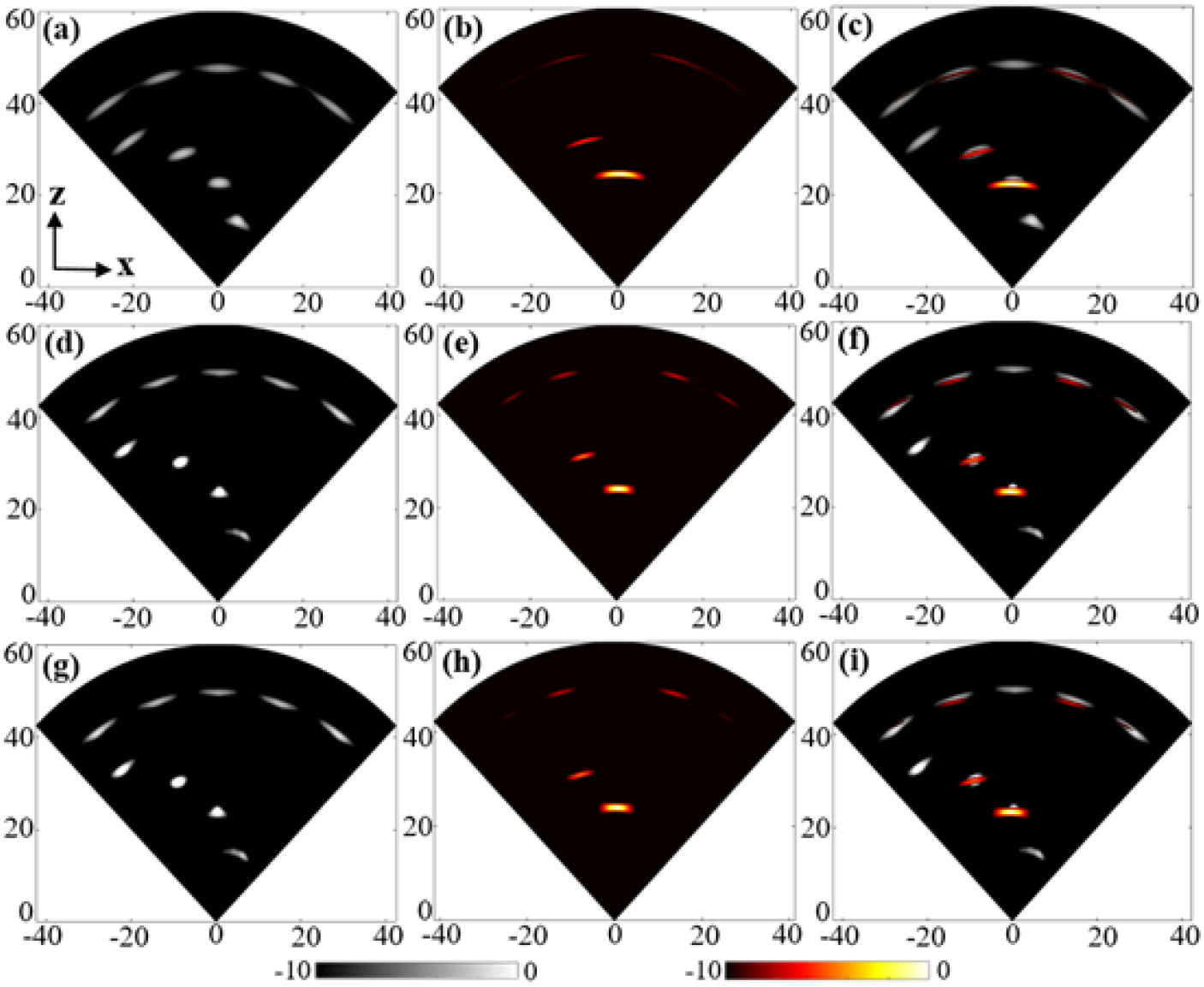
Parametric studies validating the effect of the US transducer and the light source apertures on the USPA imaging performance. Simulation results with (a-c) 40 mm light and 64-element US, (d-f) 40 mm light and 128-element US, and (g-i) 20 mm light and 128-element US array; scale: mm, colorb ar: dB.

**TABLE II:** Spatial resolution for homogeneous case studies.

| Configuration |  |  | Config.1 | Config.2 | Config.3 |
| --- | --- | --- | --- | --- | --- |
| Lateral Resolution (mm) | US | target-2 | 1.3 | 1.2 | 1.2 |
|  |  | target-8 | 1.78 | 1.72 | 1.72 |
|  | PA | target-2 | 3.52 | 1.401 | 1.44 |
|  |  | target-8 | 3.75 | 1.45 | 1.625 |
| Axial Resolution (mm) | US | target-2 | 0.57 | 0.57 | 0.57 |
|  |  | target-8 | 0.57 | 0.57 | 0.57 |
|  | PA | target-2 | 0.295 | 0.3125 | 0.32 |
|  |  | target-8 | 0.267 | 0.241 | 0.241 |

In PAI, the strength of the PA signal is directly proportional to the local optical fluence, which in turn increases with the light aperture size. Using the USPA simulation platform, we validate the effect of light aperture on the PAI performance. For a fixed ultrasound transducer array with 128-elements, when we reduced the aperture size of light illumination from 40 mm to 20 mm, the visibility of deeper photoacoustic targets (close to 5 cm) is reduced (Fig. 4e and Fig. 4h), due to the reduced optical fluence. However, no significant change in the spatial resolution was observed with the change in the size of the light source (Table II). In summary, with the increase in the size of US transducer array and the light source aperture, the visibility of both off-axis and deep tissue targets and the overall performance of both US and PA imaging improves.

### B. Effect of center frequency of ultrasound transducer

In this subsection, we study the effects of changing the center frequency of the ultrasound transducer array on the PAI performance using the tissue phantom with nine circular targets described in section II-D. The aperture sizes of light source and transducer array are fixed and the center frequency of the transducer is changed from 1 MHz, to 2 MHz and 5 MHz. The calculated lateral and axial resolutions (LR and AR in Figs. 5a-c) for the 2^nd^ target, using half the distance between 90% to 10% of the maximum PA amplitude (in the line spread functions shown in Fig. 5d and Fig. 5e), show that the spatial resolution improves with the increase in the center frequency of the US transducer array. These results confirm that the visibility of deeper targets (5 cm) become weaker with increase in the center frequency of the transducer, due to the depth and frequency dependent acoustic attenuation.

**Fig. 5.**
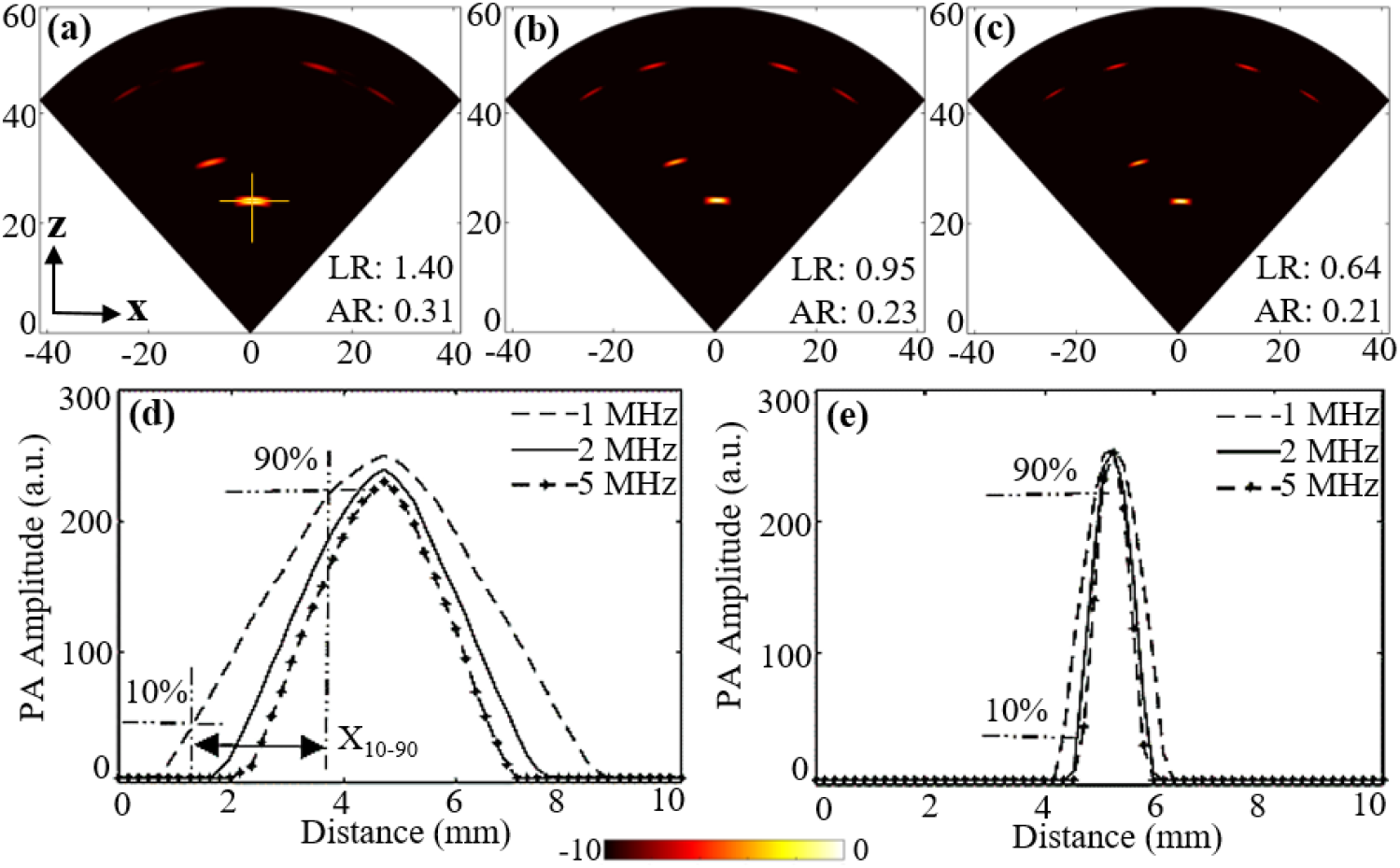
The effect of change in the center frequency of the US transducer on PAI performance. (a-c) PA images of the phantom consisting of nine-circular targets, obtained with the transducers centered at 1 MHz, 2 MHz, and 5 MHz frequency, respectively. (d, e) PA amplitudes of the 2^nd^ wire target in images (a-c), plotted in lateral and axial directions, respectively. Lateral resolution (LR) and axial resolution (AR) in mm. Scale: mm, colorbar: dB.

### C. Multispectral photoacoustic imaging and spectral unmixing

In this subsection, we present multispectral photoacoustic imaging and spectral unmixing results for the homogeneous tissue phantom embedded with HbO_2_, ICG, and Hb molecules at two different depths. Since the optical absorption of these targets is wavelength dependent, the generated pressure and resulting photoacoustic image contrast changes with the wavelength. Fig. 6a shows the position of six molecular targets: HbO_2_ (x = −15 mm), ICG (x = 0 mm), and Hb (x = 15 mm), each of radius 0.25 mm, placed at 25 mm and 40 mm depth inside the homogenous tissue background. PA images at seven wavelengths (750 nm to 900 nm, with an interval of 25 nm), were acquired using a 128-element US transducer and a 40 mm light source. Five representative PA images are shown in Figs. 6(b-f). The PA intensity plots, generated by quantifying the PA intensity of these targets in the multi-wavelength PA images (Fig. 6g), closely matches with the respective standard spectral plots of the three molecular targets (Fig. 6h). For example, the standard crossover of Hb and HbO_2_ spectra around 800 nm, and the peak response of ICG around 780 nm can be observed in these plots. We further applied a linear spectral unmixing algorithm that computes a non-negative solution to a linear least squares problem [40], using the simulated multi-wavelength PAI data, and obtained the unmixed images of HbO_2_, Hb, and ICG (Figs. 6(i-k)). These results demonstrate the feasibility of the hybrid USPA simulation platform to accurately simulate the spectroscopic PA imaging of the tissue chromophores.

**Fig. 6.**
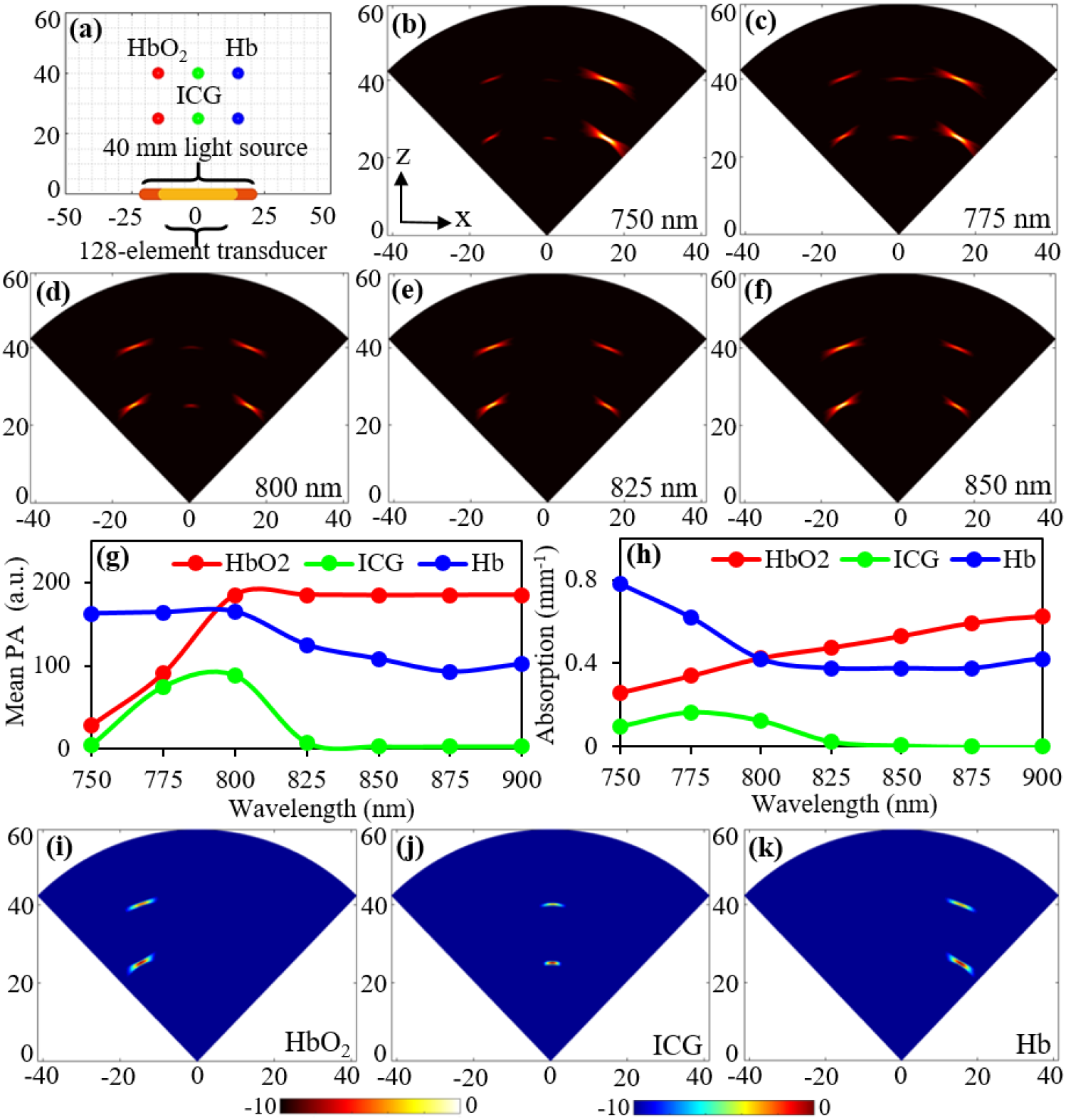
Multi-spectral PAI and spectral unmixing results with the USPA simulation platform. (a) Tissue phantom consisting of HbO_2_, ICG, and Hb molecules at two different depths. (b-f) PA images of the phantom obtained at five optical wavelengths. (g) Plots of mean PA intensities and (h) plots of optical absorption coefficients as a function of wavelength for HbO_2_, ICG, and Hb. (i-k) Corresponding spectrally unmixed images; scale: mm, colorbar: dB.

### D. Modeling USPA imaging of heterogeneous prostate tissue

The capability of our simulation platform to image heterogeneous tissue is demonstrated by developing in silico human prostate phantom and simulating TransRectal-USPA (TRUSPA) imaging of prostate using a 128-element linear US array and a 40 mm light aperture. A 600 x 600 pixels grayscale bitmap image was created with the bladder, prostate, soft-tissue and the vasculature regions. This image was converted to a 60 mm x 60 mm phantom with a gridsize of 100 μm in MatLab, as shown in Fig. 7a. The acoustic and optical properties of different tissue regions inside the prostate phantom were defined as per the literature. Figs. 7(b, c) shows the acoustic impedance and the absorption coefficient maps of the phantom. The acoustic impedances of the bladder, prostate and soft tissue were defined as 1.57, 1.60, and 1.63 respectively, in MRayls [41]. The absorption coefficient of blood vasculature, prostate, surrounding soft tissue and the bladder regions were defined as 0.425, 0.03, 0.02, and 0.01 respectively, in mm^-1^ at 800 nm [42].

**Fig. 7.**
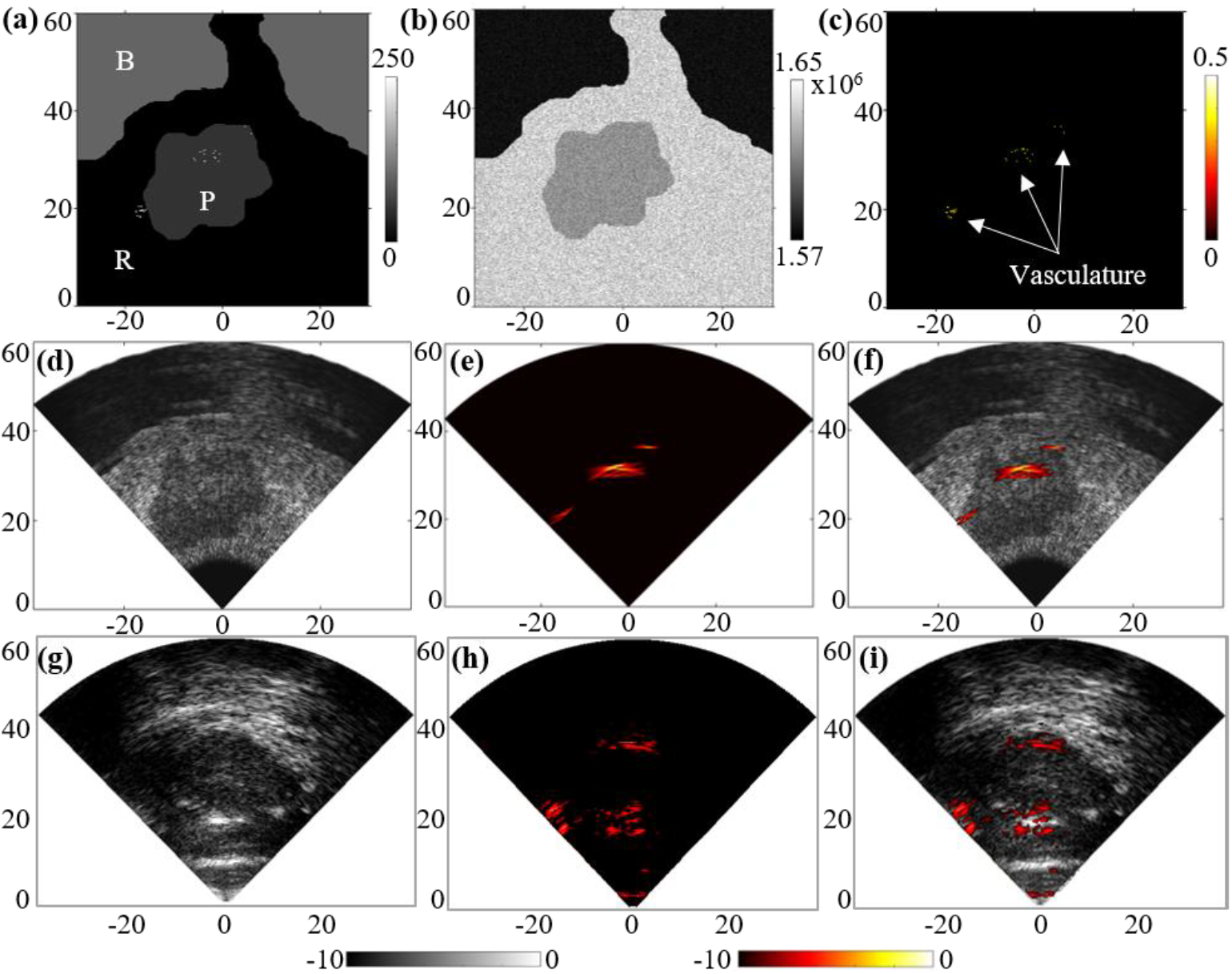
Simulating transrectal US and PA imaging of human prostate. (a) Bitmap image of the heterogeneous prostate phantom and corresponding (b) acoustic impedance map and (c) optical absorption coefficient map with arrows pointing to the prostatic vasculature. (d-f) Simulated US, PA, and coregistered US+PA images of the in silico prostate phantom. (g-i) Experimental results for in vivo TRUSPA imaging of human prostate. Bladder (B), Prostate (P) and rectal soft tissue (R). Scale: mm, colorbar: dB.

The simulated US image (Fig. 7d) clearly displays the anatomical information of the prostate, bladder (hypoechoic region above the prostate), and the surrounding tissue regions. In contrast, the simulated PA image (Fig. 7e) maps the optical absorption contrast of the prostatic vasculature. The coregistered US+PA image in Fig. 7f shows overlaid anatomical and molecular optical contrast of the in silico human prostate phantom. Figs. 7(g-i) show in vivo transrectal imaging of human prostate acquired with a TRUSPA device integrating a 64-element linear CMUT array and a fiber optic light guide, as described in section II-D [9]. These experimental results are in close agreement with the above simulation results, wherein US image shows the structure of the prostate and the surrounding regions, and the PA image shows the vasculature of the prostate and surrounding regions.

### E. USPA simulations aided deep learning enhanced PAI

Deep learning approaches are widely being investigated for medical image analysis and diagnosis, including real-time image segmentation and disease classification [43]. Despite the architectural advancements in deep learning, requirement of massive, high-quality annotated data impedes the translation of these approaches in healthcare research, where acquisition and annotation of data is seldom feasible [44].

Recently, deep learning is also being actively studied for PAI applications [45–47]. However, with PAI still in its early stage of clinical translation, the scarcity of clinical PAI data remains a major challenge in optimally training the deep learning models for a given task. Most commonly, readily available acoustic simulations have been employed for generating massive PAI datasets assuming uniform optical fluence inside the tissue medium [26], [27]. As such, these studies did not model the realistic experimental PAI where optical fluence strongly depends on the tissue depth and excitation wavelength. As demonstrated in previous sections, our model based USPA simulations account for both depth and wavelength dependent optical scattering and are capable of generating multispectral PAI datasets in realistic heterogeneous tissue environment. We present the following two studies to demonstrate the applicability of the USPA simulation datasets for deep learning enhanced PAI.

#### Application-1: USPA simulation aided deep learning approach for photoacoustic targets detection

We recently reported an encoder-decoder based convolutional neural network to identify the origin of photoacoustic wavefronts in deep-tissue scattering medium [48]. The network was trained with 16,240 model-based simulated PA images, generated by our simulation platform presented here, and tested on both simulated and experimental PAI data acquired in various background optical scattering conditions. These results demonstrated that the photoacoustic targets upto 55 mm depth can be localized with a high accuracy of ~20 μm. This can be attributed to the true modeling of optical scattering in the training PAI dataset generated by the USPA simulation platform.

Here, we demonstrate the applicability of this approach for multi-target detection by training the network with simulated PAI datasets of multiple PA targets buried in a strong background optical scattering noise. The network performance was validated using an experimental test dataset consisting of three PA targets placed inside an intralipid phantom with reduced scattering coefficient of 20 cm^-1^ (shown in Fig. 8a). Due to the heavy noise, the SNR and the visibility of deeper PA target (at 33 mm) in the conventional beamformed image is poor (Fig. 8b). However, the network trained with the PAI datasets of varying optical scattering noise conditions generated by the USPA simulation platform could precisely localize all the three PA targets (shown in Fig. 8c).

**Fig. 8.**
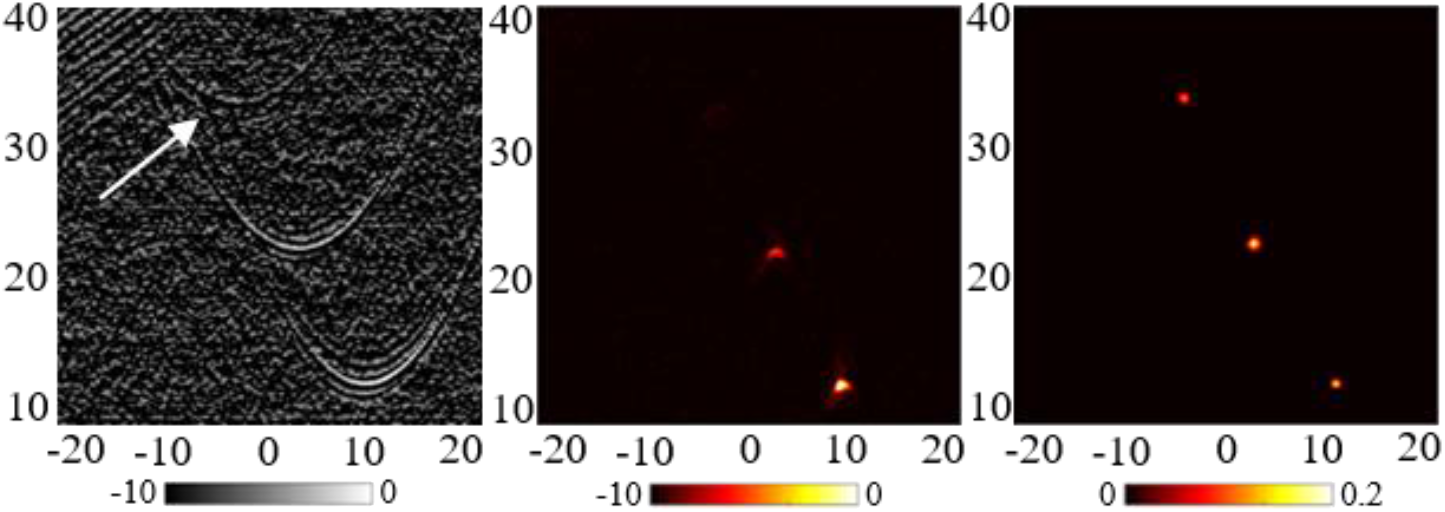
USPA simulations enabled deep-learning enhanced PAI. (a) Zoomed view of acquired raw PA data consisting of three PA targets situated at 13 mm, 23 mm, and 33 mm depth in a tissue medium with a reduced scattering coefficient of 20 cm^-1^. (b) Conventional beamformed B-mode PA image. (c) Deep-learning enhanced PA image output. Scale: mm.

#### Application-2

USPA simulations aided *unsupervised photoacoustic spectral unmixing*. Recently, we proposed an end-to-end autoencoder based neural network for delineating the molecular information in PAI [49], using the training and test multispectral PAI dataset obtained from the USPA simulation platform. These results motivated us to validate this approach for experimental PAI data in realistic tissue environments. In particular, the learned spectral features of HbO_2_ and Hb molecules from the USPA simulated training dataset (similar to Fig. 6) were used to unmix the respective molecular information present in i) a tissue phantom consisting of HbO_2_ and Hb targets (Fig. 9a) and ii) in vivo mouse prostate tumor (Fig. 9g), both acquired using the TRUSPA device described in section II-D. Figs. 9e and 9f respectively show the spectrally unmixed HbO_2_ and Hb maps obtained with the unsupervised network, when using the multispectral PAI data of tissue phantom (Figs. 9b-d) as the test input. More importantly, using the same simulated training dataset and the in vivo mouse test dataset (Figs. 9h-j), the network was able to map the HbO_2_ and Hb molecular composition inside tumor region, as shown in Figs. 9k and 9l. These results demonstrate that the multispectral PAI datasets generated by USPA simulations can adequately train the deep learning networks with the spectral knowledge of tissue chromophores and subsequently help unmix the molecular information in realistic in vivo environments.

**Fig. 9.**
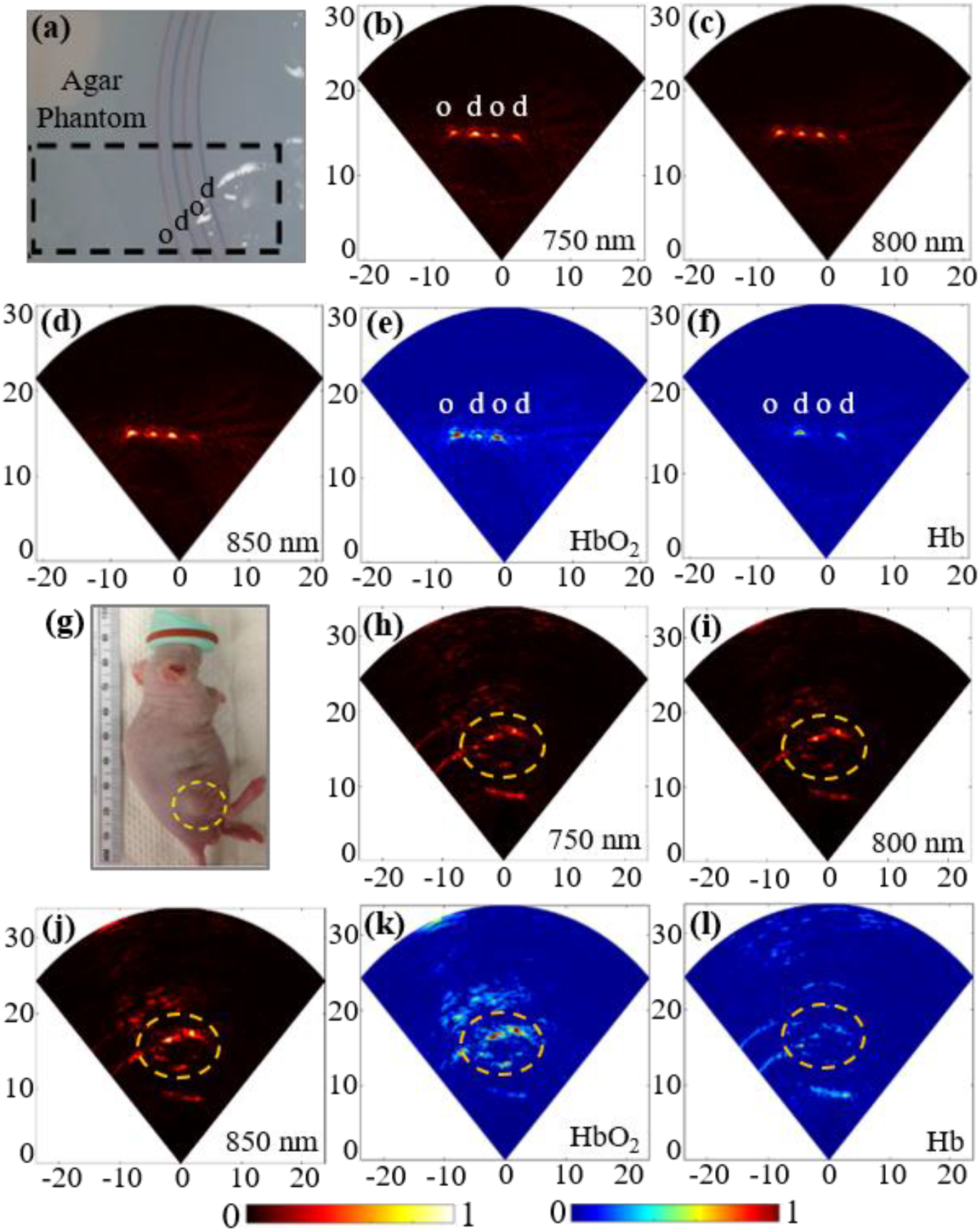
USPA simulation aided unsupervised PA spectral unmixing for (a) a tissue mimicking phantom with tubes filled with HbO_2_ ‘o’ and Hb ‘d’. (b-d) PA images of the phantom acquired at three wavelengths. (e, f) Corresponding unmixed results for HbO_2_ and Hb. (g) In vivo mouse tumor. (h-j) PA images of the mouse tumor acquired at three wavelengths. (k, l) Corresponding unmixed results for HbO_2_ and Hb. Scale: mm.

## IV. Conclusion

This paper presented and validated a hybrid USPA numerical simulation approach that adequately simulates multispectral PAI as well as US imaging of deep (upto 60 mm) homogeneous and heterogeneous biological tissue. The simulation platform integrates two open source toolboxes, i) NIRFast for forward light propagation and calculation of optical fluence at each tissue grid location; and ii) K-Wave for ultrasound propagation and detection. The platform models dual-modality USPA devices and generates B-mode US images featuring anatomical information, multispectral PA images displaying functional and molecular information based on optical absorption contrast, and a combined US+PA image with overlaid anatomical and molecular contrasts. Extensive parametric studies demonstrated that the USPA simulations can practically model the effect of key design parameters such as the size of the US transducer array, the light source aperture, and the frequency of the US transducer on the USPA imaging performance. In addition, the capabilities of the USPA simulation platform to accurately map the spectral profiles of deep tissue molecular targets and obtain the respective unmixed molecular information has been demonstrated. Furthermore, the feasibility of USPA imaging of heterogeneous tissue was demonstrated by i) developing an in silico human prostate phantom mimicking the acoustic and optical properties of the prostate tissue; and ii) generating simulated transrectal US (anatomy of the prostate and surrounding regions), PA (prostatic vascular contrast) and US+PA images of the prostate.

In addition to assisting the device modeling, this paper also presented the applicability of using the USPA simulated training datasets in aiding deep-learning enhanced PAI. This was tested over experimental datasets for i) localizing deep-tissue vascular targets buried in strong optical scattering noise and ii) delineating the HbO_2_ and Hb molecular information inside a phantom as well as in vivo mouse tumor.

The presented USPA simulations platform provides a powerful tool for optimizing the performance of USPA imaging devices for clinical applications. More importantly, the heterogeneous tissue modeling and imaging capabilities of the simulation tool opens the door for several emerging deep-learning applications of PAI by supplying sufficient realistic training datasets. Future scope of this work involves 3-D simulation and validation studies on diverse organs (beyond prostate demonstrated in this work) mimicking realistic optical and acoustic heterogeneities, artifacts, shadow effects and system noise.

## Acknowledgement

This work was supported in part by the NIH-NIBIB R00EB017729-04 and by Penn State Cancer Institute startup funds. We also acknowledge the support of NVIDIA Corporation with the donation of Titan X Pascal GPU.

